# YAP1 Regulates the Self-organized Fate Patterning of hESCs-Derived Gastruloids

**DOI:** 10.1101/2021.03.12.434631

**Authors:** Servando Giraldez, Eleonora Stronati, Ling Huang, Hui-Ting Hsu, Elizabeth Abraham, Kathy A. Jones, Conchi Estaras

## Abstract

During gastrulation, the coordinated activity of BMP, WNT and NODAL signaling pathways guide the differentiation of the pluripotent epiblast into the three germinal layers. Recent studies underline the role of the Hippo-effector YAP1 regulating WNT and NODAL signaling pathways and repressing mesoendodermal differentiation in human embryonic stem cells (hESCs). However, the contribution of YAP1 to the cell-fate patterning decisions that transform the epiblast in a three-germ layer gastrula remains unknown. We address this question by analyzing micropatterned 2D-gastruloids derived from hESCs, in the presence and absence of YAP1. Our findings show that YAP1 is necessary for gastrulation. YAP1 KO-gastruloids display reduced ectoderm layer and enlarged mesoderm and endoderm layers, compared to WT. Furthermore, YAP1 regulates the self-organized patterning of the hESCs, as the discrete position of the three germ layers is altered in the YAP1 KO-gastruloids. Our epigenome (single-nuclei ATACseq) and transcriptome (RNA-seq) analysis revealed that YAP1 directly represses the chromatin accessibility and transcription of key genes in the NODAL pathway, including the NODAL and FOXH1 genes. In WT gastruloids, a gradient of NODAL: SMAD2.3 signaling from the periphery to the center of the colony regulates the exit of pluripotency toward endoderm, mesoderm and ectoderm, respectively. Hence, in the absence of YAP1, a hyperactive NODAL signaling retains SMAD2.3 in the nuclei impeding the self-organized differentiation of hESCs. Accordingly, the partial inhibition of NODAL signaling is sufficient to rescue the differentiation and pattern -defective phenotypes of the YAP1 KO gastruloids. Our work revealed that YAP1 is a master regulator of NODAL signaling, essential to instruct germ layer fate patterning in human gastruloids.

## INTRODUCTION

Gastrulation is a crucial developmental stage when the pluripotent epithelial epiblast subsequently converts into the three embryonic germ layers: mesoderm, ectoderm, and endoderm^1^. The success of the gastrulation largely relies on the correct spatial and temporal regulation of the NODAL pathway^2-4^. NODAL:SMAD2.3 signaling is essential for maintaining pluripotency in pre-gastrulating epiblast cells^5-8^. However, as development proceeds, NODAL cooperates with BMP and WNT pathways in the posterior epiblast to initiate Primitive Streak (PS) formation, the precursor of mesoderm and endoderm layers^1^. Convergently, inhibitory signals of NODAL and BMP in the anterior epiblast facilitate the acquisition of a neuroectodermal potential^1, 9^. Thus, epiblast cells interpret a high-medium-low level of NODAL signal as a “sign” to either integrate into the PS, remain pluripotent, or differentiate into a neuroectodermal cell, respectively. Yet, the underlying mechanisms that control NODAL signaling are not entirely understood.

NODAL pathway ligands are members of the transforming growth factor-beta (TGFβ) superfamily that bind to type I and type II serine-threonine kinase receptors. Activated type I receptors phosphorylate cytoplasmic SMAD2 and SMAD3, leading to their interaction with Smad4 and the subsequent formation of transcriptional complexes in the nucleus^2^. The antero-posterior gradient of NODAL:SMAD2.3 signaling during gastrulation is partially shaped by inhibitory extraembryonic signals from the anterior visceral endoderm (AVE), including Lefty1 and Cer1^10,11^. However, epiblast-like cells (hESCs and mESC) in vitro organize in absence of extraembryonic tissues^12-14^; in 2D-gastruloids, the germ layers organize in a defined concentric pattern in confined hESC colonies^12^, and hESC-derived 3D-gastruloids break symmetry and display anteroposterior organization^13, 14^. These observations emphasize the relevance of intrinsic mechanisms in epiblast cells regulating the spatial distribution of NODAL cues, independent of polarized extraembryonic signals.

The Hippo pathway regulates organ size, regeneration, and cell growth by controlling the stability of the transcription factors YAP1 and TAZ. Hippo/MST kinase cascades phosphorylate YAP1/TAZ, leading to their ubiquitin-mediated degradation. When the Hippo kinases are inactive, YAP1/TAZ translocate into the nucleus where they associate with TEAD1–4 DNA-binding proteins and regulate transcription^15-17^. Unlike TAZ, YAP1 is essential for development, and YAP1 KO mice die around E8.5^18^. The YAP1 KO mouse showed defects in extraembryonic and embryonic tissues, including axis elongation^18^. Lineage-specific YAP1 KO mutants have revealed specific roles of YAP1 in the cell-fate decisions of the zygote^19^ and specification of the trophectoderm and epiblast-cells^20-22^. Similarly, we and others have described important roles of YAP1 during differentiation of epiblast-like cells (hESCs and mESCs)^23-28^. For instance, YAP1 represses mesoendodermal (ME) differentiation in hESCs^24, 25, 29^ by regulating the activity of the NODAL and WNT3 pathways^24, 26^. However, whether YAP1 regulates gastrulation signaling during the formation and spatial organization of the three-germ layers has not been addressed.

Here, we used a robust model of human gastrulation^30, 31^ to examine the YAP1-dependent regulatory mechanisms. In this model, hESCs are confined to circular micropatterns and differentiated following exposure to BMP4 cytokine. Our findings indicate that YAP1 is required for normal BMP4-induced gastruloid formation. YAP1 KO-derived gastruloids exhibit expansion of mesoderm and endoderm layers and substantial reduction of ectoderm-specified cells. Moreover, our data show that YAP1 controls the patterning of the gastruloids, as the YAP1 KO-gastruloids display abnormal location of the three-germinal layers, compared to WT. Our genome-wide analysis reveals that the predominant role of YAP1 in human gastrulation is to repress the expression of regulatory genes of the NODAL:SMAD2.3 pathway. Accordingly, the gastrulation-defective patterning in YAP1 KO hESCs can be partially rescued by inhibiting NODAL signaling. Overall, our data highlights YAP1’s essential role in controlling embryonic spatial patterning and underlying cell-intrinsic regulatory mechanisms in hESCs.

## RESULTS

### YAP1 is required for cell-fate patterning of hESC-derived 2D gastruloids

To examine the role of YAP1 in human gastrulation, we analyzed the differentiation pattern of the three-germinal layers in 2D-gastruloids derived from hESCs^12^. For this, the hESCs were grown in geometrically confined discs (circular CYTOO chips) until they reached ∼80-90% confluency. Then, micropatterned colonies were treated with BMP4 cytokine to differentiate them into self-organized concentric rings of embryonic germ layers; with ectoderm in the center, extra-embryonic tissue at the edge, and mesoderm and endoderm in between^12^ (**Figure 1A, WT**). We compared the patterned structures derived from WT and YAP1 KO hESCs. We found that the number of ectoderm positive cells, expressing the SOX2 marker^32^, is significantly reduced in the YAP1 KO gastruloids, compared to WT. However, the percentage of cells positive for mesoderm (Brachyury/T^+^) and endoderm (SOX17^+^) markers was higher in the YAP1-deleted micropatterns (**Figure 1A, 1B and Supplemental Figure1**). Consequently, YAP1 KO gastruloids display expanded mesoderm and endoderm rings, and reduced ectodermal ring, compared to WT (**Figure 1A, 1D and Supplemental Figure1)**. Furthermore, the distribution of the concentric rings is inverted in the YAP1 KO gastruloids, and the ectoderm cells are located in the periphery, surrounding the mesoderm cells, that now occupy the center of the 2D-gastruloid (**Figure 1A, 1C, 1E and Supplemental Figure 1**). Altogether, these findings reveal that YAP1 regulates the fate and allocation of the three germinal layers in hESCs.

**Figure 1.**
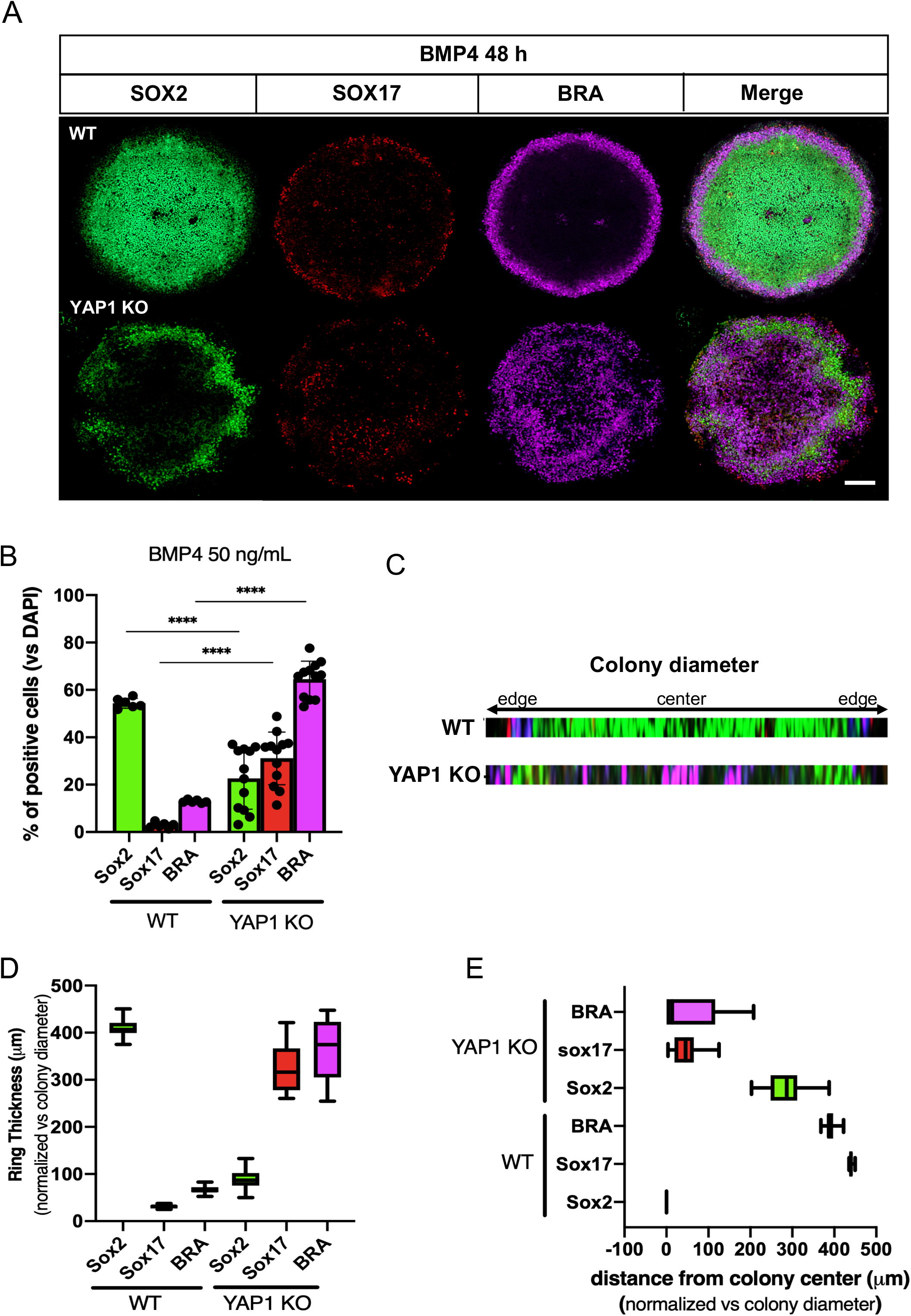
YAP1 regulates the formation of the three germinal layers in hESC-derived 2D-gastruloids. **A)** Immunostaining for the germinal layer markers SOX2 (Ectoderm), SOX17 (Endoderm) and Brachyury (Mesoderm) in WT and YAP1 KO micropatterned cell cultures stimulated with BMP4 (50 ng/mL) for 48 hrs. **B)** Percentage of cells positive for SOX2, SOX17, and Brachyury markers in WT and YAP1 KO 2D-gastruloids, versus the total number of cells in the pattern. **C)** Cross-section showing the distribution profile of the germ-layer markers. **D-E)** Graphs show quantification of ring thickness (D) and distance from colony center (E) of WT and YAP1 KO-derived gastruloids. Experiments were replicated three times with consistent results. 10-15 colonies were analyzed. Scale bar 100 µm. Statistical significance was quantified by applying unpaired two-tailed t-test. P values ****≤0.001.

### YAP1 KO hESCs fail to differentiate into ectoderm

The analysis of the 2D-gastruloids suggest that the acquisition of an ectodermal cell-fate is compromised in the YAP1-depleted cells. To better examine the contribution of YAP1 to the formation of each of the germinal layers, we used a directed differentiation protocol (StemCell Technologies #05230) to induce differentiation towards ectoderm, mesoderm, and endoderm lineages in WT and YAP1 KO hESCs (**Figure 2A**). After culturing the cells for 5 days (mesoderm and endoderm) or 7 days (ectoderm) with the lineage specific medium, we examined the transcriptome of the differentiated cells using RNA-seq approaches. The heatmap in **Figure 2B** shows the expected germ-layer markers following the induction of the three lineages in WT hESCs. However, the YAP1 KO cells failed to express ectoderm lineage genes (**Figure 2B**). Instead, after 7 days of exposure to ectodermal-induction media, the YAP1 KO cells expressed high levels of pluripotency markers, such as POU5F1 and NANOG, and low levels of key ectodermal genes, including SOX1 and PAX6 (**Figure 2B, 2C**). Importantly, similar to what we and others have previously shown, the induction of endoderm genes was enhanced in the YAP1-KO cells compared to WT^24-26,29^ (**Figure 2B, 2C)**. Previous reports have shown that NANOG represses ectoderm differentiation but has little effect on other lineages^33^. Accordingly, in the WT hESCs, NANOG expression decreases in the first 24h of ectoderm differentiation, preceding the induction of the ectoderm gene, OTX2 (**Figure 2D**). However, the YAP1 KO hESCs display standing levels of NANOG following 72h of ectoderm induction, which correlates with the absence of ectoderm genes (**Figure 2B, 2D**). We restored the levels of YAP1 in the YAP1 KO hESCs using a Doxycycline-inducible PiggyBac system^24^. Our data show that the reintroduction of YAP1 is sufficient to reduce the levels of NANOG and induce OTX2 expression in ectoderm induced-YAP1 KO hESCs (**Figure 2E-F**). Altogether, these findings show that YAP1 is necessary to repress NANOG during hESC-directed differentiation toward an ectodermal fate.

**Figure 2.**
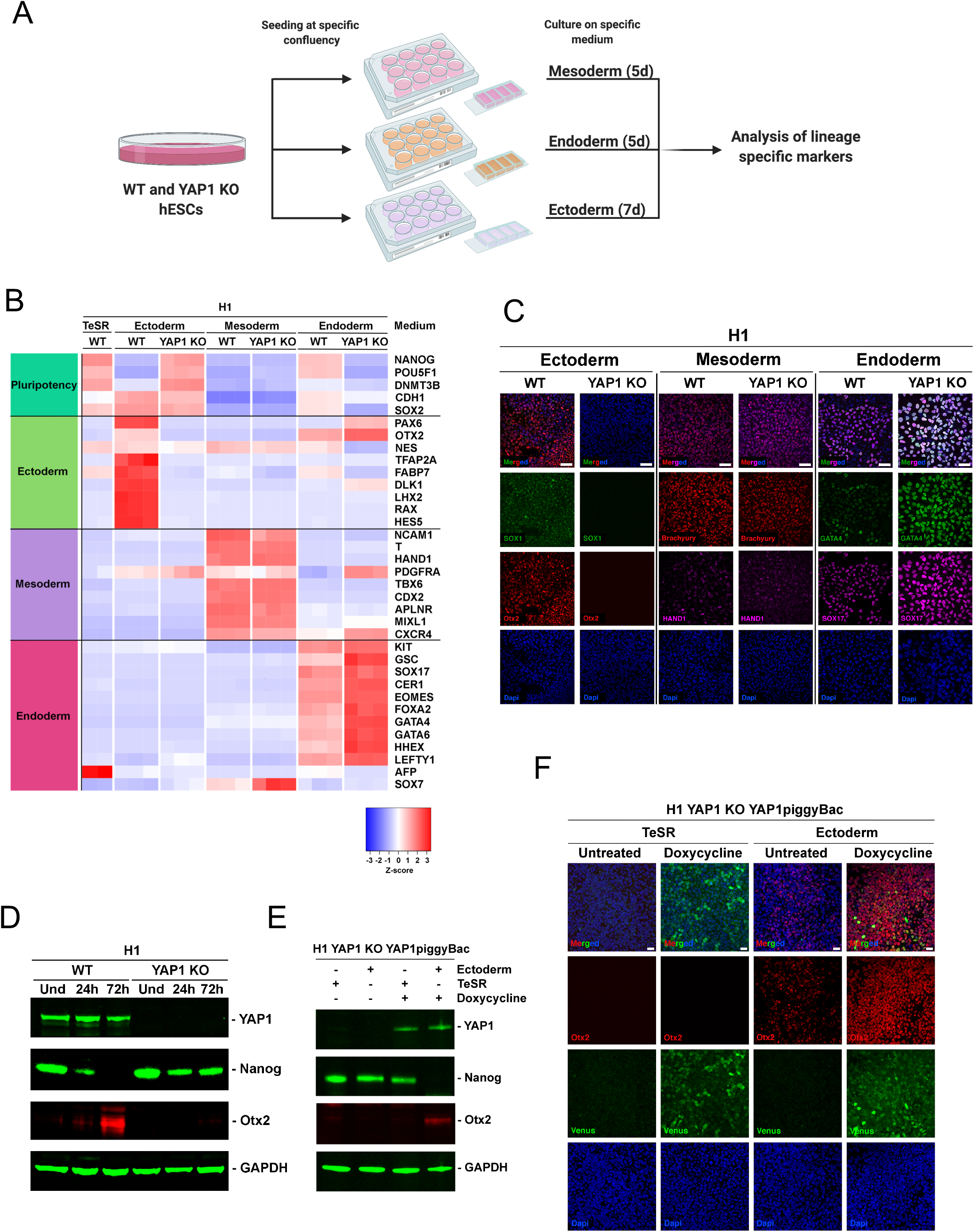
YAP1 is essential for the ectoderm fate commitment after directed differentiation. **A)** Schematic diagram of the directed differentiation methodology. **B)** Heat map from RNA-seq analysis showing expression levels of pluripotency and lineage specific genes following directed differentiation using the conditions shown in 2A. (n=3, Z scaled log2 FPKM). The cell lines and treatments are indicated above the map. **C)** Immunostaining of WT and YAP1 KO cells following the directed differentiation methodology shown in 2A. The cells are stained against specific lineage markers; Ectoderm: SOX1 (Green) and Otx2 (Red); Mesoderm: Brachyury (Red) and HAND1 (purple); Endoderm: GATA4 (Green) and SOX17 (Purple). Nuclei are stained in DAPI. Scale bar 50 µm, (n=3). **D)** Representative immunoblot of the indicated proteins in WT and YAP1 KO cells grown on mTeSR1 (undifferentiated, Und) or ectoderm medium for 24 or 72 hours, as shown. GAPDH was used as a loading control, (n=3). **E)** Representative immunoblot of the indicated proteins. The YAP1 KO:YAP1PiggyBac stable cell line was generated by transfection of the transposon-based vector PiggyBac containing an intact Flag-YAP1-Venus cDNA under a Doxycycline promoter. Cells were grown on mTeSR1 or ectoderm medium for 72 hours and treated with 20 ng/ml doxycycline, as indicated, (n=3). **F)** Immunostaining of YAP1 KO:YAP1PiggyBac cells after the same treatment described above. The immunostaining shows the levels of an ectoderm specific marker Otx2 (red) and the expression of the doxycycline-regulated Flag-YAP1-Venus (green). Nuclei are stained in DAPI. Scale bar 25 mm, (n=3).

### Cytoplasmic retention of SMAD2.3 during ectoderm induction it depends on YAP1

TGFb/NODAL:SMAD2.3 signaling maintain the expression of pluripotency genes, including NANOG^5^. Accordingly, dual inhibition of TGFb/NODAL:SMAD2.3 and BMP:SMAD1.5 signals is sufficient to turn off the pluripotency program and promote the acquisition of an ectodermal fate in hESCs, similar to the in vivo situation^9, 34^. We interrogated whether YAP1 regulates SMADs during ectoderm induction in hESCs. For this purpose, we examined the cellular localization of the SMAD2.3 and SMAD1.5 in WT and YAP1 KO cells. The subcellular fractionation experiments in **Figure 3A** show that global SMAD2.3 levels are higher in the YAP1-KO cells than in the WT cells. More importantly, the SMAD2.3 failed to translocate into the cytoplasm in response to ectoderm induction media in the YAP1 KO cells (**Figure 3A and Supplemental Figure 2A, 2B)**. Interestingly, we did not detect differences in the cytoplasmic accumulation of SMAD1.5 in the YAP1-KO cells (**Supplemental Figure 2A, 2B**). Then, we examined SMAD2.3 activity in an *in vitro* model of 2D-neurulation in WT and YAP1 KO hESCs. We seeded the hESCs in geometrically confined colonies and treated them with “homemade” neurulation media, which contains the TGFb/NODAL and BMP inhibitors: SB431542 and Noggin, as previously described^35^. After 4 days of neurulation induction, SMAD2.3 was vastly located in the cytoplasm of the WT cells. However, the SMAD2.3 remained predominantly nuclear in the YAP1 KO colonies (**Figure 3B**), consistent with the lack of expression of the ectoderm marker SOX2 (**Figure 3B**). Overall, these findings show that YAP1 is necessary to inhibit the nuclear activity of SMAD2.3 during the commitment toward neuroectodermal fates.

**Figure 3.**
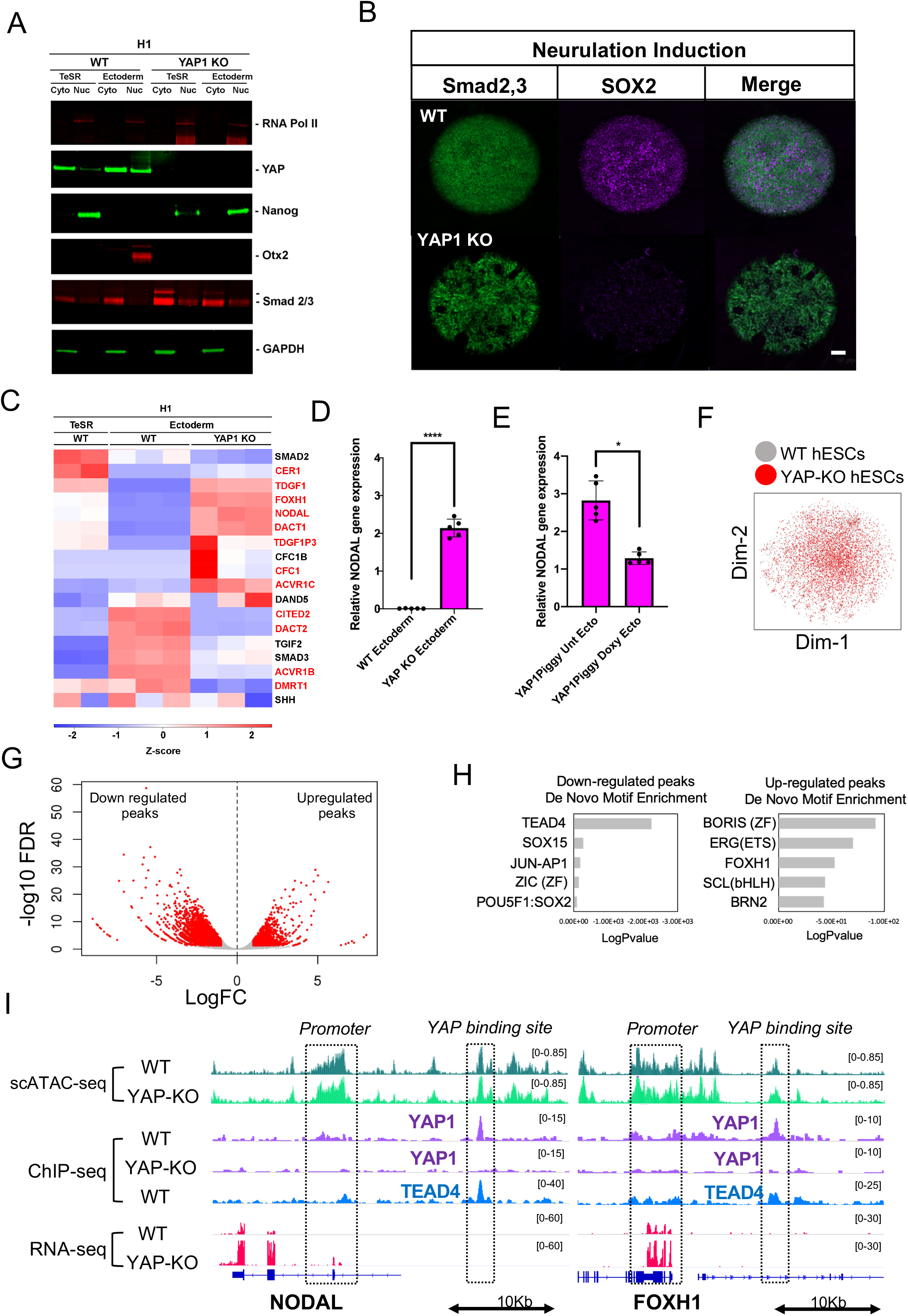
YAP1 inhibits NODAL:SMAD2.3 signaling during ectodermal differentiation induction. **A)** Cells were grown on mTeSR1 or ectoderm medium for 72 hours. Representative immunoblot of the indicated proteins in WT and YAP1 KO cells after Cyto/nuclear sub-fractioning followed by Western Blotting analysis (n=3). **B)** Immunostaining of indicated markers in WT and YAP1 KO-derived neuruloids differentiated on 500 µm diameter CYTOO chips following 4 days in neurulation induction media. Note the nuclear retention of SMAD2.3 (green) and the reduced expression of the ectodermal SOX2 (purple) in YAP1 KO, compared to WT. Scale bar 50 µm. **C)** Heatmap from RNA-seq experiment in Figure 2 depicts NODAL pathway regulatory genes (AmiGo database) in WT and YAP1 KO hESCs differentiated to the ectoderm. For comparison, self-renewing hESCs are also shown (mTeSR1, WT). The genes differentially regulated in WT and YAP1 KO cells are highlighted in red. Colors represent z-scaled log2-transformed FPKM values. **D-E)** Graphs show qPCR analysis of NODAL mRNA levels in the WT and YAP1 KO cells (D) or YAP1 KO:YAP1PiggyBac cells in the presence or absence of doxycycline (E).Note that restoring YAP1 levels (+doxy) is sufficient to reduce the mRNA levels of NODAL. **F-I)** Analysis of snATAC-seq experiments in WT and YAP1 KO hESCs. **F)** t-distributed stochastic neighbor embedding (t-SNE) showing unbiased clustering results for all the sequenced cells. The plots are colored by sample identity (WT hESCs:grey, YAP1 KO hESCs: red) **G)** Volcano plot shows differentially regulated (DR) regions in YAP1 KO hESCs (2,462 peaks upregulated and 8,221 downregulated at false discovery rate < 0.05 log2 fold-change > 1). The plots are colored by significance (red = significant, grey = not significant). **H)** sc-ATACseq analysis of DNA motifs differentially enriched in the YAP1 KO versus WT hESCs **I)** Genome browser captures of snATAC-seq (UCSC), RNA-seq and ChIP-seq of the indicated proteins, in WT and YAP1 KO hESCs. Note that there is a correlation between increase chromatin accessibility and mRNA levels for NODAL and FOXH1 genes, in the YAP1 KO versus WT cells. ChIP-seq identified a YAP1 binding site in enhancers near NODAL and FOXH1 genes.

### YAP1 represses NODAL pathway genes during ectoderm differentiation

We hypothesized that the nuclear retention of SMAD2.3 was due to an abnormal NODAL signaling in the YAP1 KO gastruloids. To test this idea, we examined the expression levels of regulatory components of the NODAL signaling pathway in ectoderm-induced WT and YAP1 KO cells. For that, we applied a Signaling Pathway Analysis to the RNA-seq datasets derived from the WT and YAP1 KO cells after ectoderm induction (**Figure 2A**). The AmiGO database^36^ classified 18 genes in the NODAL pathway (GO0038092). We identified 12 of the genes differentially regulated (DR) in the YAP1 KO cells compared to WT (**Figure 3C**, DR genes are highlighted in red, overlapping p-value < 0.001, Fisher’s Exact Test), suggesting that the NODAL pathway is strongly affected by YAP1 deletion. The list of DR genes contains important NODAL signaling regulators, such as the transcription factor FOXH1 (Log2 Fold Change 4.69, p-value < 0.001), an essential partner of SMAD2.3 in gastrulation^37^. We also identified upstream signaling components strongly upregulated in the YAP1 KO cells, such as the Activin receptor ACVR1C (log2 fold-change 1.92, adjusted p-value < 0.001) and the NODAL gene (log2 fold-change 13.42, adjusted p-value < 0.001) (**Figure 3C, D**). Furthermore, restoring YAP1 levels using the Doxycycline-inducible PiggyBac system^24^ in the YAP1 KO cells was sufficient to reduce the NODAL gene expression during ectodermal induction (**Figure 3E**). These results suggest that YAP1 represses the transcription of NODAL signaling genes during ectodermal differentiation.

### YAP1 regulates the chromatin accessibility of FOXH1 and NODAL developmental genes

To better understand the transcriptional regulatory mechanisms of YAP1 in gastrulation, we applied the cutting-edge single-nuclei ATAC-seq (snATAC-seq) technology to examine the genome-wide chromatin accessibility profile in WT and YAP1-KO hESCs. The t-distributed stochastic neighbor embedding (t-SNE) analysis in **Figure 3F** revealed that WT and YAP1 KO hESCs clustered together in a homogenous cell population. These findings indicate that the overall chromatin profiling of YAP1 KO hESCs resembles normal pluripotent cells. However, we identified 10,683 differentially regulated (DR) regions in the YAP1 KO cells compared to WT (**Figure 3G**). We classified the 10,683 DR peaks into proximal (+/-10Kb from TSS) and distal (>10Kb from TSS) genic regions. This analysis revealed that 8,767 out of 10,683 DR regions lie in the distal regions, and most of them lost accessibility in the YAP1 KO hESCs (83%). This result suggests that YAP1 facilitates enhancer accessibility, in agreement with the previous reports^38^. However, this analysis also revealed that up to 49.8% of the proximal regions (promoters) gained accessibility in the YAP1 KO cells (954 out of 1917 DR promoters) (**Supplemental Figure 2C**). Thus, YAP1 has important roles in restricting accessibility on promoters. As expected, the DNA motif most represented in the regions that lost accessibility in the absence of YAP1 was TEAD4, followed by SOX15 and JUN-AP1 motifs (**Figure 3H)**. Convergently, the DNA motifs recognized by BORIS, ERG, and FOXH1 transcription factors gained access in the YAP1 KO cells. Importantly, among the regions that gained accessibility in the YAP1 KO hESCs, we identified the promoter of NODAL and FOXH1 genes **(Figure 3I**), which correlates with an increase in the mRNA levels of these genes (**Figure 3C-3E and 3I**). Furthermore, our ChIP-seq analysis of YAP1 in WT and YAP1 KO hESCs^24^ revealed two bona-fide YAP1-binding sites upstream of the NODAL and FOXH1 genes (**Figure 3I**), suggesting a direct role of YAP1 repressing these genes. The dynamic regulation of the NODAL gene during embryogenesis is orchestrated by the activity of cell-type specific enhancers^11^ (**Supplemental Figure 2D**). We identified that the YAP1-bound region near the NODAL locus corresponds to the well-characterized Proximal Epiblast Enhancer (PEE). The PEE drives NODAL expression in the epiblast and Primitive Streak in vivo, but it is inactive in the anterior epiblast^11, 39^. Our data suggest that YAP1 represses the PEE to inactivate the NODAL gene during ectoderm induction. These observations are in agreement with previous reports describing a transcriptional repressor role of YAP1 on developmental genes^24, 26, 40, 41^. Finally, our genome comparison analysis revealed that YAP1:TEAD binding site in the PEE is conserved in mouse (**Supplemental Figure 2D**), suggesting a preserved role of YAP1 regulating the NODAL enhancer. Overall, we conclude that YAP1 restricts the chromatin accessibility of the NODAL and FOXH1 genes by regulating key developmental enhancers.

### Partial inhibition of the NODAL pathway rescues the phenotype of YAP1 KO gastruloids

Our data show that YAP1 represses the NODAL and FOXH1 genes to restrain the SMAD2.3 nuclear activity during ectoderm induction. Thus, we hypothesized that the defective gastrulation phenotypes of YAP1 KO hESCs are due to the persistent activity of NODAL:SMAD2.3 signaling throughout the gastruloid. To test this idea, we examined the effect of blocking NODAL signaling in the YAP1 KO-gastruloids. Hence, we replicated the 2D-gastrulation experiments in the YAP1 KO cells, in the presence and absence of the NODAL receptor inhibitor A83-0159^42^. **Figure 4** shows that the co-treatment of BMP4 and the NODAL inhibitor (**Figure 4A**) rescues the expression and patterning of the ectoderm layer (SOX2+) in the YAP1 KO 2D-gastruloids, compared to BMP4 treatment alone (**Figure 4A-D**). Accordingly, an outer layer of Brachyury+ cells is visualized around the SOX2+ cells in the YAP1 KO gastruloids exposed to BMP4+A83-0159, partially resembling the original structure of WT gastruloids (**Figure 4A-D**). We noticed that longer exposures to A83-0159 (48h) completely blocks the activity of SMADs, and even compromised the formation of the mesoderm and endoderm layers (**Supplemental Figure 3A, 3B**). Finally, we tested whether the A83-0159 was sufficient to rescue the YAP1 KO phenotypes during ectoderm directed differentiation (**Figure 2A**). For this, WT and YAP1-KO hESCs were exposed to commercial ectoderm induction media for 3 days in the presence and absence of A83-0159. As shown in **Supplemental Figure 3C and 3D**, blocking NODAL signaling was sufficient to downregulate NANOG and induce the expression of the ectoderm lineage gene OTX2 in the YAP1 KO cells. Altogether, these findings highlight the importance of YAP1 regulating NODAL signaling to coordinate cell-fate decisions during human gastrulation.

**Figure 4.**
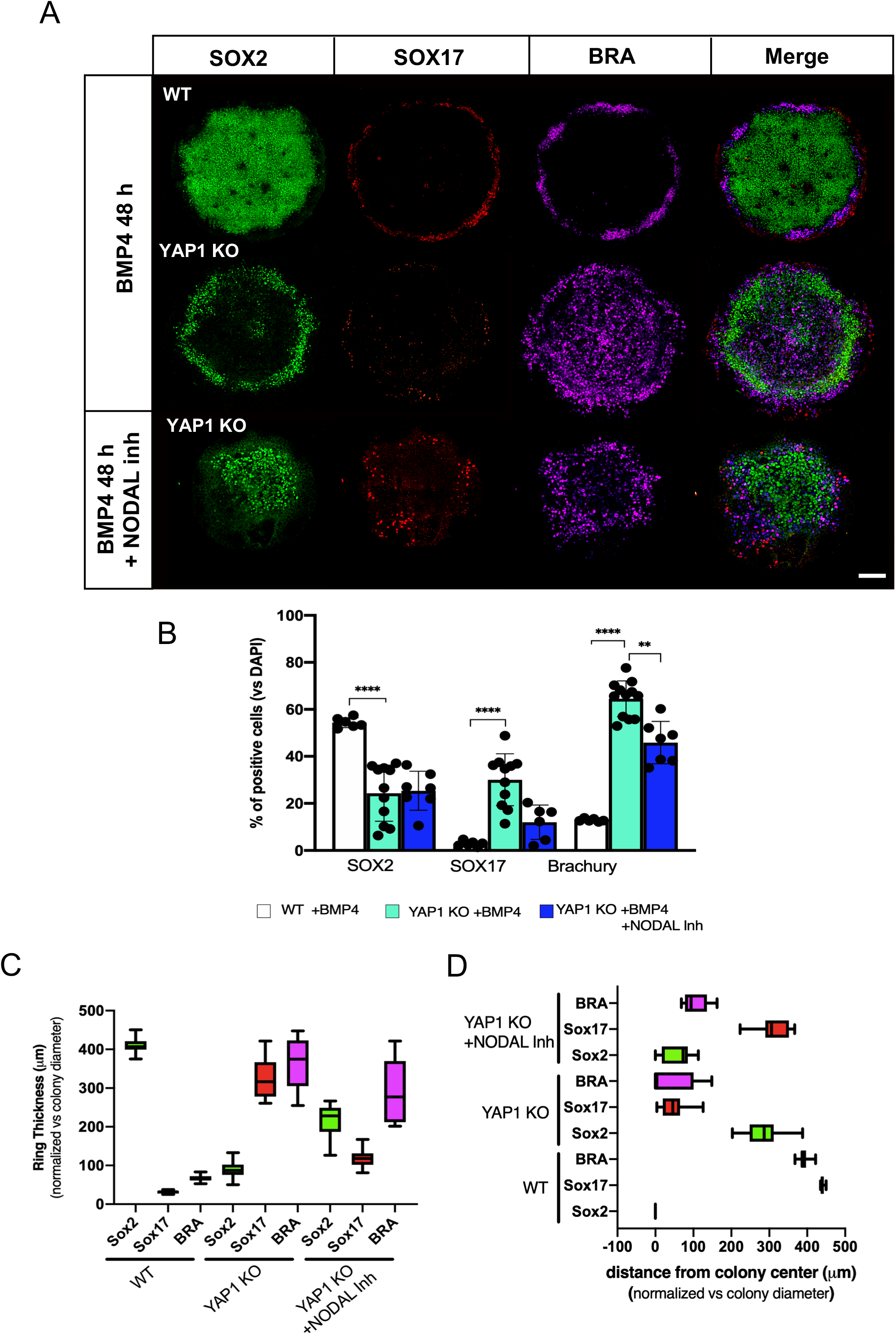
Nodal inhibition rescues the 2D-gastrulation defects in YAP1 KO cells. **A)** Immunostaining for the germinal layer markers SOX2 (Ectoderm), SOX17 (Endoderm) and Brachyury (Mesoderm) in the YAP1 KO micropatterned cell cultures stimulated with BMP4 (50 ng/mL) or combined BMP4+ Nodal Inhibitor (A8301 1mM). WT hESC-derived gastruloids are shown for comparison. **(B)** Graph shows percentage of SOX2+, SOX17+ and Brachyury+ cells of WT and YAP1 KO in the presence or absence of Nodal Inhibitor. **C-D)** Graphs show quantification of ring thickness (**C)** and distance from colony center (**D**). 10-15 colonies were analyzed. Scale bar 100 µm. p value **≤0.01, ****≤0.001, unpaired two-tailed t-test.

## CONCLUSIONS

Directed differentiation strategies in hESCs have previously shown that YAP1 regulates the expression of lineage genes during the specification of the mesoderm and endoderm fates^24-26,29^. However, the process of gastrulation requires additional coordinated actions of epiblast cells to self-organize and choose the correct differentiation path toward one of the three germ-layers. Thus, we interrogated the role of YAP1 in the process of gastrulation in hESCs. Our studies revealed three main findings: 1) YAP1 regulates the correct specification and allocation of the three-germ layers in the 2D-gastruloids, 2) YAP1 is needed for the induction of an ectodermal fate in hESCs, and 3) a predominant role of YAP1 repressing the activity of NODAL signaling. Importantly, mutations in NODAL signaling ^43-46^ or YAP1^47-50^ have been identified in Holoprosencephaly and Colobomas. Holoprosencephaly and Colobomas belong to the same spectrum of developmental diseases^51-53^ and are suspected to originate during gastrulation^51, 54^, by abnormal patterning of anterior neuroectoderm^51^. Our findings established a novel connection between NODAL and YAP1 pathways in gastrulation, opening the door to future studies on investigating the NODAL:YAP1 regulatory axis in the context of abnormal development and birth defects.

## MATERIAL AND METHODS

### hESC culture and cell lines

Control, YAP1 KO and the doxycycline-inducible YAP1 (YAP1 KO:FlagYAP1-PiggyBac) H1 hESCs were previously described^24^. hESCs were cultured in mTeSR1 medium on Matrigel-coated tissue culture plates. For expansion, hESC colonies were fragmented and split in a 1:20 ratio following ReLeSR treatment (StemCell Technologies). For directed differentiation experiments, colonies were disaggregated into single cells using Accutase and plated in the presence of Rock Inhibitor (StemCell Technologies Y-27632) on matrigel-coated plates. The YAP1 KO:FlagYAP1-PiggyBAC cells were treated with 20 ng/ml of Doxycycline to restore the YAP1 expression. For 2D-gastrulation experiments see Methods in Micropatterns Cell Culture section.

### Directed differentiation

hESCs were single-cell seeded onto 6-well Matrigel-coated plates at 2*10^6^ cells/well for ectoderm and endoderm conditions and 5*10^5^ cells/well for mesoderm condition and treated with the appropriate differentiation medium using the STEMdiff Trilineage Differentiation Kit (StemCell Technologies #05230) according to manufacturer instructions. Differentiated cells were then collected for RNA isolation, fixed with 4% paraformaldehyde for immunostaining or harvested for western blotting analysis.

### Micropattern cell culture and Gastrulation induction

Micropatterned glass chips from CYTOO (CYTOOCHIP Arena 1000, France) were used, following Deglincerti et al. protocol^32^. Briefly, the chips were coated with 10 mg/mL of Cell-Adhere Laminin-521 (StemCell Technologies) diluted in PBS with calcium and magnesium (PBS++) (Sigma) for 2h, at 37°C. Before seeding the cells, chips were washed with PBS++. hESCs were dissociated with Accutase, pelleted in DMEM/F12 (Gibco) and resuspended in growth media. 1*10^6^ cells in 2.5 mL were seeded on the chips with Rock Inhibitor. After 4h, Rock Inhibitor was removed and replaced with standard mTeSR1 media for 12-16 hrs. Cells were then washed with PBS and treated with 50 ng/mL of BMP4 supplemented with PenStrep (Sigma) for 48hrs.

### Neurulation induction

For neurulation experiments cells were seeded in mTeSR1 for 12-16h then washed twice with PBS before adding neurulation media (DMEM-F12, 15% KnockOut Serum Replacement media (Gibco), 0.2% NEAA (Sigma), 500 ng/mL Noggin (R&D #6057-NG/CF), 10 mM SB431542 (Stemgent #04-0010-10)) supplemented with PenStrep (Sigma). This media was freshly prepared and replaced daily until collection on day 4.

### Treatments with the Activin/NODAL/TGF-β pathway inhibitor

Cells were grown on TeSR1 or ectoderm medium and treated with the Activin/NODAL/TGF-β pathway inhibitor 2mM A83-0159 (StemCell Technologies #72022) for 72h. In 2D-gastrulation experiments the chips were stimulated with BMP4 for 48h and 1mM of A83-0159 was added to the media with BMP4 (48h treatment) or after 12 hrs (36h treatment).

### Immunostaining, imaging, and quantification

For direct differentiation experiments, hESCs were plated in Lab-Tek II Chambered Coverglass slides previously coated with Matrigel. Cells were fixed with formaldehyde 4% for 10 minutes. After permeabilization with Triton 0.1% for 10 minutes, cells were washed with PBS, and incubated with blocking solution (PBS Tween 0.1%, BSA 0.1%, FBS 10%) for 30 minutes at room temperature. Fluorophore-conjugated primary antibodies (See **Supplemental Table 1**) diluted in blocking solution were added and incubated overnight at 4°C. After washes, cells were incubated with 1µg/ml DAPI for 10 minutes. Finally, cells were washed three times with PBS. Images were captured by a Nikon Eclipse Ti confocal microscope using Nikon NIS-Element AR software. For immunostaining of micropatterned gastruloids, CYTOO chips were washed twice with PBS before fixing with freshly prepared 4 % formaldehyde (Life Technologies) for 20 minutes at room temperature. The chips were then washed twice with PBS, and the cells were permeabilized with blocking solution (PBS 0.1% Tryton-X, 3% of Donkey Serum (Jackson Immunoresearch Cat#017-000-121)) for 45 minutes at RT. Primary antibodies (listen in **Supplemental Table 1**) were added overnight at 4°C diluted to the appropriate concentration in blocking solution. The following day, the chips were washed three times in PBS 0.1% Tween-20 (PBS-T) for 15 minutes, followed by incubation with secondary antibodies in the presence of 1 mg/mL of DAPI (Invitrogen) in blocking solution for 1h, at RT. Finally, chips were washed three times in PBS-T for 15 minutes at room temperature and one final wash in PBS. The chips were mounted on glass slides with mounting media (DAKO, Cat#S3023). Micropatterned gastruloids were imaged using Zeiss Airyscan Confocal LSM900; tile Z-stack images of 5-6 colonies were collected using 20x objective. Each channel was saved as a separate tiff file and analyzed with Fiji software. All analysis was carried out in Fiji, homogenously subtracting the background from tile images. Each colony was then analyzed using single-channel images. Cell nuclei were quantified using the analyze particle plugin in Fiji and expressed in a graph as a percentage of positive cells compared to the DAPI staining. The distance from colony center and ring thickness were manually quantified, averaging 4 repeated measurements for each colony. Otherwise stated, an average of 10-15 colonies were quantified for each experiment.

### RNA extraction and qPCR

RNA was extracted using Quick RNA Zymo kit following manufacturer indications. 0.5 µg of total RNA was reverse transcribed using Transcriptor First Strand Synthesis kit (Roche). The cDNA was amplified using SYBR green master mix (Life Technologies) on a CFX96 Touch Real-Time PCR detection system. All results were normalized to RPS23, GAPDH and Actin genes. The ΔΔCt method was used to calculate relative transcript abundance against the indicated references. Unless otherwise stated, error bars denote standard deviation among three to five independent experiments.

### RNA-sequencing and bioinformatic analysis

WT and YAP1 KO hESCs were differentiated into ectoderm, mesoderm, or ectoderm layers, following directed differentiation strategies (see directed differentiation section in Methods). RNA was extracted using Zymo columns and sent for library preparation and quality control to the Salk Institute Genomics Analysis Laboratory (SIGnAL). High-throughput sequencing was carried out in the Illumina HiSeq 4000 device. Raw reads were mapped to the human hg19 reference by STAR aligner (v2.5.3a)^55^ using default parameters. Then reads uniquely mapped across exons of RefSeq genes were summed by HOMER (v4.9.1)^56^ to be used as gene expression counts. Differential regulated (DR) genes were identified using DESeq2 (v1.18.1)^57^ with adjusted p-val < 0.01 and absolute log2 fold-change (log2FC) > 1. Fragments per kilobase per million mapped reads (FPKM) values were log2-transformed and z-scaled for heatmap visualization. A pseudo-value of 5 was added before log2-transformation. Heatmap was generated using R package “gplots” [ref: https://cran.r-project.org/web/packages/gplots/index.html].

### Single-nuclei ATAC-sequencing (snATAC-seq) and bioinformatic analysis

Sample preparation, library generation and single-cell sequencing snATAC-seq was carried out at the Center for Epigenomics at UCSD. The procedure followed the protocol described in Cusanovich et al^58^. Briefly, DMSO-frozen hESCs were thawed in 37°C water bath until liquid (∼2-3 minutes) and immediately placed on ice. Nuclei were isolated and permeabilized using OMNI buffer (0.1% NP40, 0.1% Tween-20, 0.01% Digitonin) as described^59^. Nuclei were counted and 2k nuclei were dispensed per well; 96 wells per sample for tagmentation. After tagmentation (and addition of the first barcode), nuclei from individual samples were pooled. 20 nuclei from each sample were sorted per well (768 wells total). Following nuclear lysis the second barcode was added, and DNA molecules were pooled and purified. Before sequencing, the quality of the library was certified by analyzing fragment size distribution and DNA concentration. Sequencing was performed in a HiSeq 4000 device.

SnapTools (v1.4.7) and SnapATAC (v1.0.0) developed at Ren’s Lab, UCSD were used for the scATAC-seq analysis^60^. In brief, reads were mapped to hg19 reference using bwa mem (v0.7.12)^61^ and sorted by samtools (v1.9)^62^. Then reads were pre-processed by SnapTools using default parameters and chromosome bin size of 5000. Cells with less than 500 unique molecular identifier (UMI) reads were removed from downstream analysis. The distribution of counts in promoter ratio was inspected and cells with the ratio < 0.1 or > 0.6 were removed. MACS2 (v2.1.2)^63^ was used for peak calling from pooled reads (parameters: -g hs --nomodel --shift 37 --ext 73 --qval 1e-2 –B --SPMR --call-summits). Chromosomal bins overlapped with the ENCODE hg19 blacklist regions, random genomic contigs, or mitochondrial chromosomes were removed from analysis. Top 5% most expressed bins were also removed. Dimension reduction by Latent semantic analysis (LSA) was performed on binary chromosomal bins and visualized by t-distributed stochastic neighbor embedding (tSNE) using the top 30 principle components and a random seed of 10. Differential regulated (DR) regions were called by SnapATAC command “findDAR” using parameters “bcv=0.1, fdr=0.05, pvalue=0.01, test.method=exactTest, seed.use=10”. DR regions with false discovery rate (FDR) < 0.05 and absolute log2 fold-change > 1 were considered as significantly different. Motifs enrichment analysis was performed by HOMER (v4.9.1)(http://homer.ucsd.edu/homer/), with parameters “-size 200 -mask”.

### Western Blotting

Cells were harvested and total cell extracts were prepared in cold lysis buffer (50 mM Tris pH 8.0, 150 mM NaCl, 1 mM EDTA, 1% Triton X-100, 1 mM PMSF, 10 mM NaF, 1 mM Na3VO4, 2 µg/mL aprotinin, 1 µg/mL leupeptin, 1 µg/mL pepstatin), sonicated, and cleared by centrifugation for 10 minutes at 10,000 g at 4C. Protein concentration was determined by Bradford protein assay (Bio-Rad), sample buffer was added (50 mM Tris pH 6.8, 2% SDS, 0.01% bromophenol blue, 2.5% beta-mercaptoethanol and 10% glycerol), and extracts were incubated for 5 minutes at 100C. Samples were separated in 10% SDS-PAGE gels (Invitrogen) and transferred onto PVDF membranes using iBlot2 gel transfer device (ThermoFisher). Membranes were blocked for 1 hour at room temperature in Super Block™ (ThermoFisher) blocking buffer, incubated overnight at 4C with primary antibody (See table X) in blocking buffer, and then incubated for 1 hour at room temperature with secondary antibody in blocking buffer. The protein bands were visualized using the Odyssey Infrared Imaging System (LI-COR Biosciences, Lincoln, NE, USA).

### Cytosolic/nuclear fractions

hESCs were subjected to subcellular fractionation after directed differentiation using the Cell Fraction Kit (Cell signaling Technology #9038S). The cytosolic and nuclear extracts were prepared following manufacturer instructions to be analyzed by western blotting as described above.

### Accession numbers

Full data sets of scATACseq (from WT and YAP1 KO H1 hESCs) and RNAseq (ectoderm, mesoderm and endoderm cells derived WT and YAP1 KO hESC) have been submitted to the NCBI Gene Expression Omnibus database in a MIAME-compliant format under superseries accession number GSE168377. The ChIP-seq of YAP1 and TEAD4 in hESCs can be found in <u>GSE99202</u>.

## AUTHOR CONTRIBUTIONS

CE and HS conceived the project. CE supervised and designed the experiments, wrote the manuscript and attained funding. KAJ supervised directed differentiation experiments and attained funding. SG designed and performed directed differentiation experiments and analysis, and prepare figures. ES designed and performed experiments in the micropatterns, their corresponding analysis, and prepared figures. EA helped with the micropattern experiments and analysis of the PEE. SG, ES, HS and EA reviewed and edited the manuscript. LH performed all bioinformatics analysis of the sequencing data.

## ACKNOWLEDGMENTS AND FUNDING

We acknowledge support from Salk Institute Bioinformatics and Stem Cell Cores. We also acknowledge support from the Center for Epigenomics at the University of California, San Diego (UCSD). This work is funded by the Temple Seed Fund and WW. Smith Charitable Foundation grant (to CE) and by the California Institute for Regenerative Medicine (<u>GC1R-06673-B</u>) (to KAJ).

Human models of 2D gastruloids require

