## SUPPLEMENTAL FILES for "YAP1 Regulates the Self-organized Fate Patterning of hESCs-Derived Gastruloids"

Supplemental Figure 1

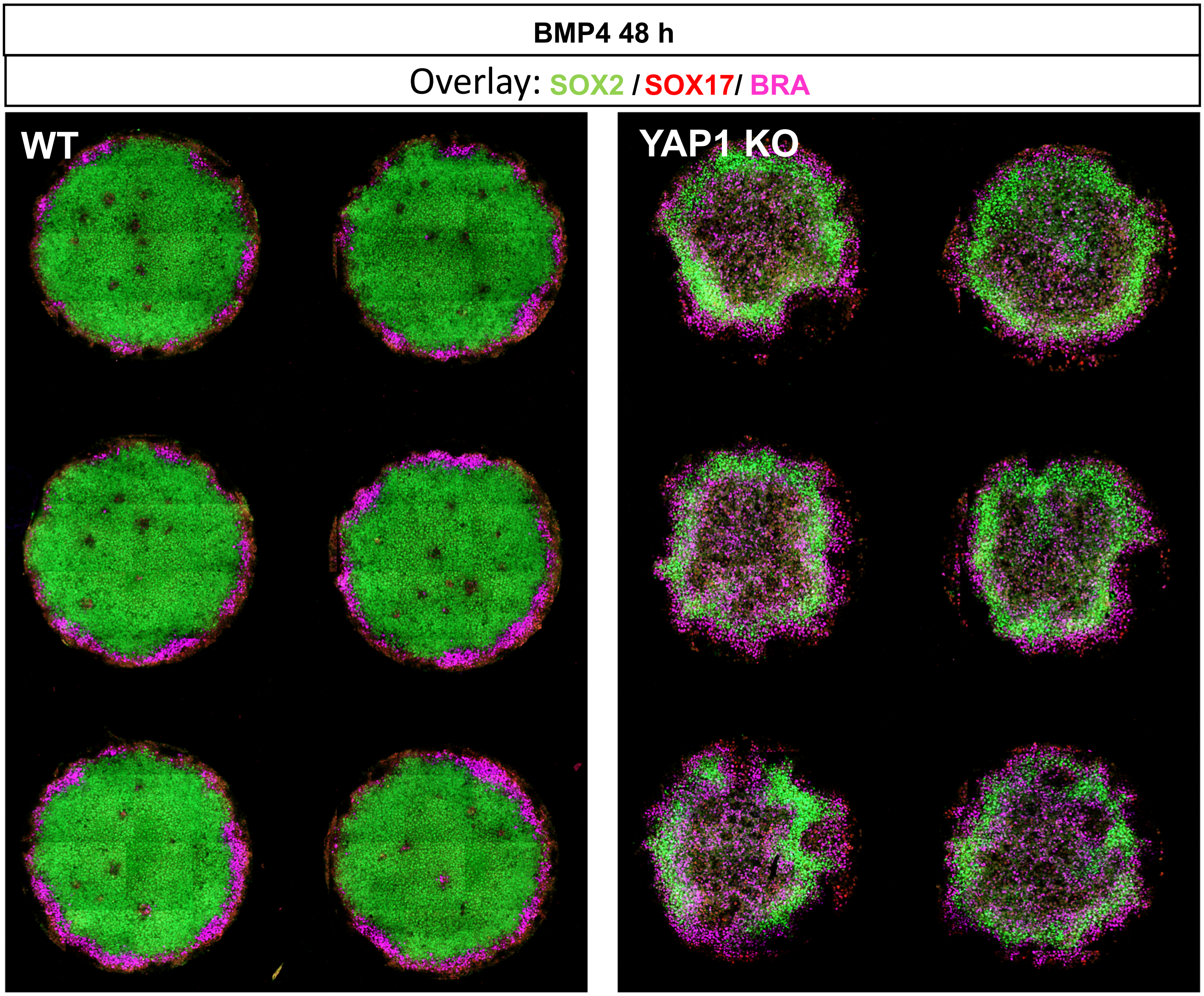

**Supplemental Figure 1.** Tile images of micropatterned WT and YAP1 KO gastruloids, stimulated with BMP4 (50ng/mL) for 48 hrs. Immunostaining show markers for Ectoderm (SOX2; green), Mesoderm (Brachyury; purple), and Endoderm (SOX17; red).

### Supplemental Figure 2

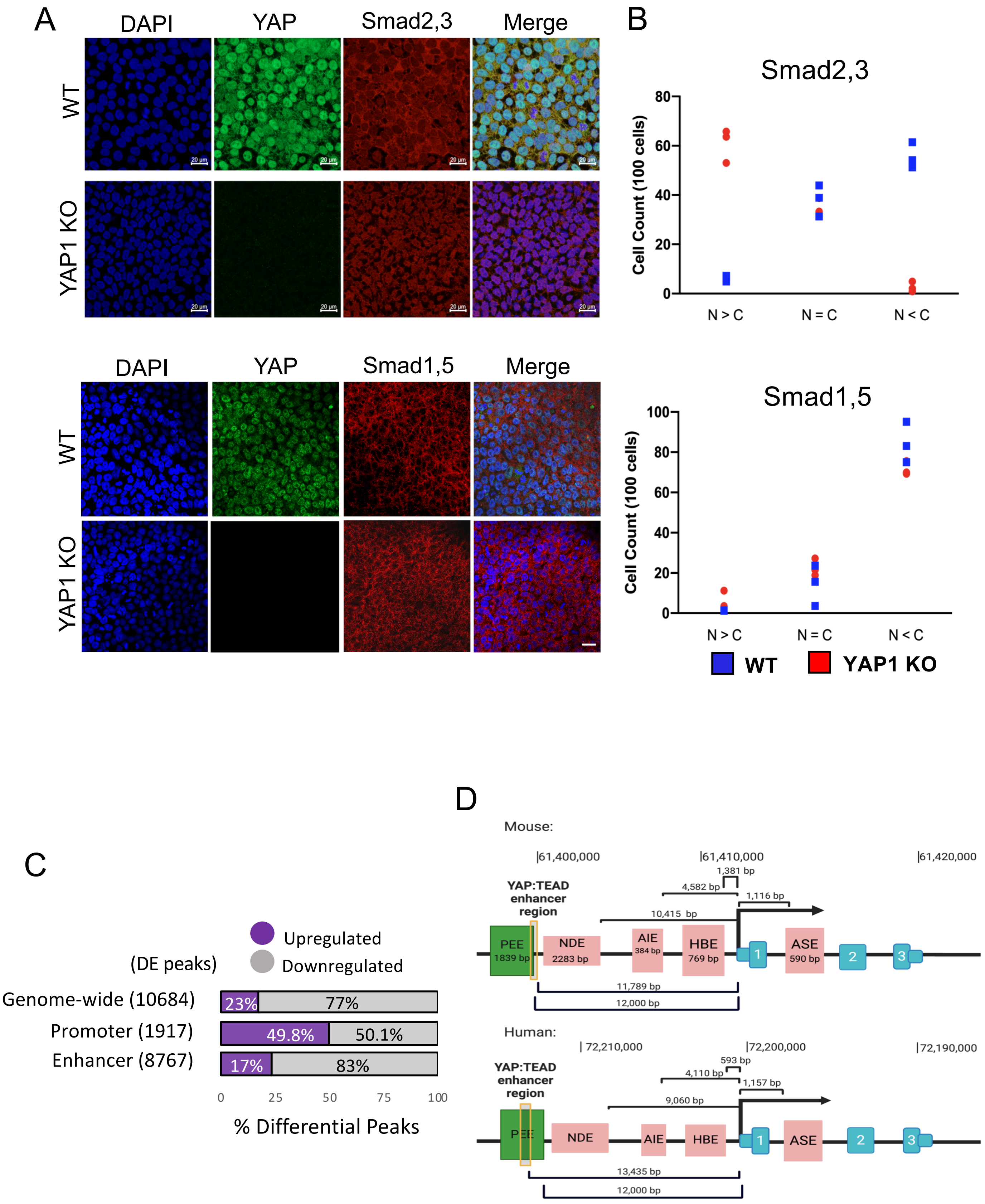

**Supplemental Figure 2. A)** Immunostaining for SMAD2.3 (upper panel) and SMAD1.5 (lower panel) in WT and YAP1 KO after 2 days of ectodermal induction. **B)** Quantification of subcellular localization of Smads in WT and YAP1 KO cells. N>C: mainly nuclear localization; N=C: equally distributed between nucleus and cytoplasm; N<C: mainly cytoplasmic localization. Scale bar 50 mm. (100 cells were counted per condition). **C)** Scheme from scATAC-seq analysis shows differential expressed peaks in YAP1 KO versus WT hESCs (genome-wide; 10,684). The scheme also displays upregulated and downregulated peaks classified by distance to the nearest TSS (promoter: <10Kb, enhancer:>10Kb). **D)** Scheme of the NODAL gene for humans and mice with characterized cis-regulatory enhancers; Proximal Epiblast Enhancer (PEE), Nodal node-specific Enhancer (NDE), Asymmetric Initiator Element (AIE), Highly Bound Element (HBE), and Asymmetric Element (ASE). The enhancer distances to the TSS are indicated, and the blue boxes represent the NODAL exons.

Supplemental Figure 3

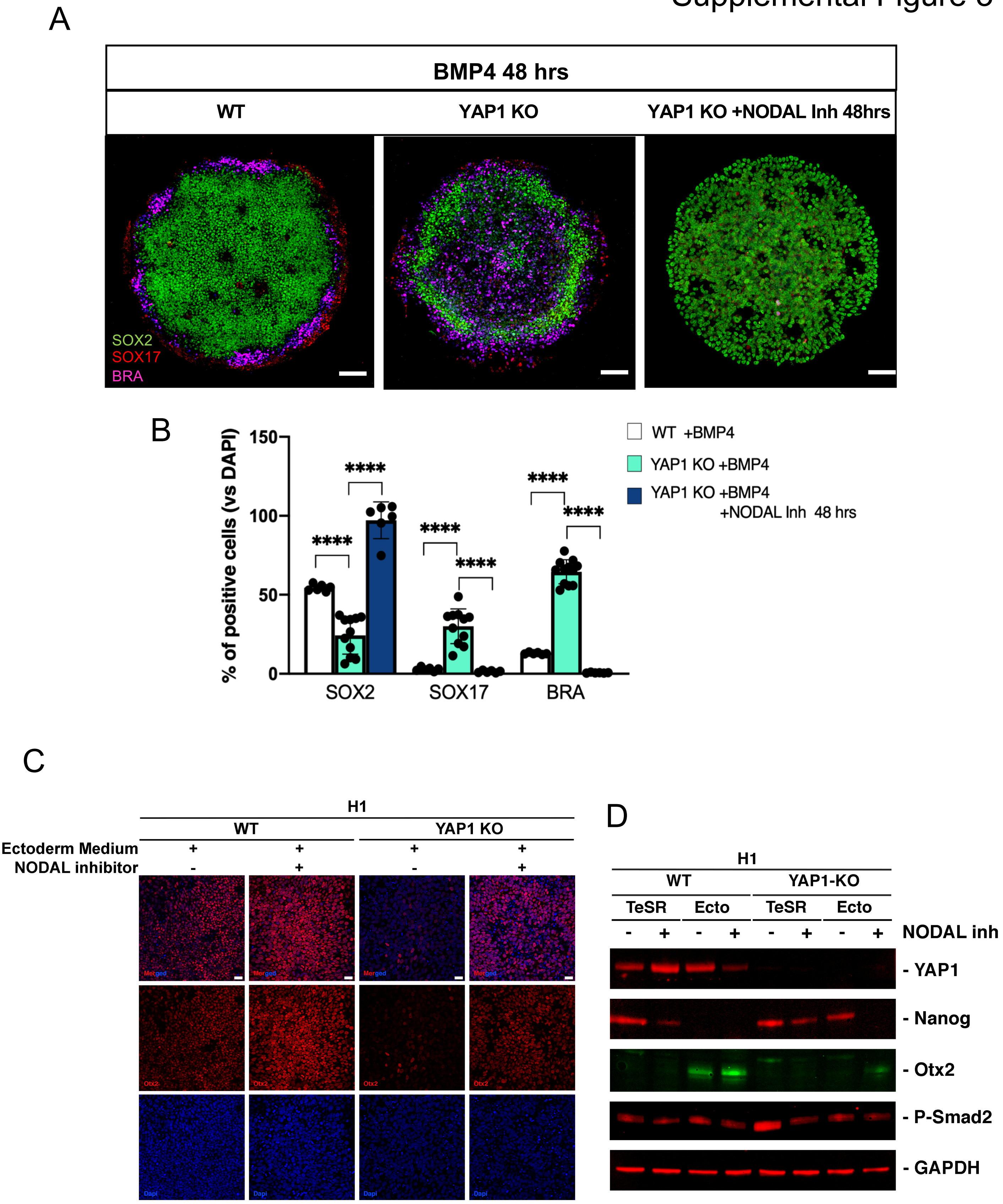

**Supplemental Figure 3. A)** Immunostaining for SOX2 (green), SOX17 (red) and Brachyury (purple) of YAP1 KO cells treated with BMP4 (50 ng/mL) and NODAL inhibitor (A83-0159 1mM) for 48h. Note that A83-0159 treatment was added together with BMP4, for a total of 48h. In the main Figure 4, the A83-0159 was added 12h later, for a total of 36h. Scale bar 100mm (BMP4-treatment images), and 50mm (BMP4+A83-0159 images). **B)** Percentage of SOX2, SOX17 and Brachyury-positive cells in YAP1 KO-gastruloids treated with BMP4, in the presence or absence of NODAL inhibitor, for 48 h. p values ****< 0.001 (unpaired Student t-test). **C)** Immunostaining showing OTX2 levels in WT and YAP1 KO hESCs following ectoderm directed differentiation for 72h, in the presence or absence of the NODAL inhibitor (A83-0159, 2 mM). Nuclei are stained in DAPI. Scale bar 25 mm. **D)** WT and YAP1 KO hESCs were grown on mTeSR1 or ectoderm medium for 72 hours, in the presence or absence of the NODAL inhibitor (A83-0159, 2mM). Representative immunoblot of the indicated proteins from total cell extracts is shown (n=3).

Table S1

| Primary antibodies | Sp. | Company/ source | Catalog | Use |
|---|---|---|---|---|
| anti-GAPDH | Rabbit | Cell signaling | 5174S | Western Blotting |
| anti-YAP1 | Rabbit | Santa Cruz | sc-15407 | Western Blotting |
| anti-Nanog | Rabbit | Cell signaling | 4903S | Western Blotting |
| anti-Otx2 | Goat | RnD | AF1979 | Western Blotting/IF |
| anti-RNA Pol II | Rabbit | Santa Cruz | sc-899 | Western Blotting |
| anti-Smad 2/3 | Rabbit | Santa Cruz | sc-83332 | Western Blotting |
| anti-SOX1 NL493-Cojugated | Goat | RnD | 967390 | Immunofluorescence |
| anti-Otx2 NL557-Conjugated | Goat | RnD | 967389 | Immunofluorescence |
| anti-Brachyury NL557-Conjugated | Goat | RnD | 967388 | Immunofluorescence |
| anti-HAND1 NL637-Conjugated | Goat | RnD | 967392 | Immunofluorescence |
| anti-GATA-4 NL493-Conjugated | Goat | RnD | 967391 | Immunofluorescence |
| anti-SOX17 NL637-Conjugated | Goat | RnD | 967393 | Immunofluorescence |
| anti-Sox2 | Mouse | RnD | MAB2018 | Micropattern |
| anti-Sox2 | Goat | RnD | AF2018 | Micropattern |
| anti-Brachyury | Goat | RnD | AF2085 | Micropattern |
| anti-Sox17 | Mouse | Abcam | ab3310 | Micropattern |
| anti-Smad 2/3 | Mouse | Santa Cruz | sc-133098 | Micropattern |
| anti-Smad1,5,8 | Rabbit | Santa Cruz | sc-6031-R | Micropattern |
| **Secondary antibodies** | **Company/ source** | **Catalog** | **Use** | |
| Goat anti-Rabbit Cross-Absorbed Dylight 800 | Invitrogen | SA5-10036 | Western Blotting | |
| Donkey anti-Goat IRDye 800CW | LI-COR | 925-32214 | Western Blotting | |
| Donkey anti-Goat Highly Cross-Adsorbed Alexa Fluor Plus 680 | Invitrogen | A32860 | Western Blotting | |
| Alexa Fluor® 488 | Jackson Immunoreasearch | | Immunofluorescence | |
| Alexa Fluor® 594 | Jackson Immunoreasearch | | Immunofluorescence | |
| Alexa Fluor® 647 | Jackson Immunoreasearch | | Immunofluorescence | |
| **qPCR genes** | **Forward** | **Reverse** | | |
| NODAL | GAGATTTTCCACCAGCCAAA | AGGTGACCTGGGACAAAGTG | | |
| RPS23 | TGTCGTGGACTTCGTACTGC | ATGCCACTTCTGGTCTCGTC | | |
| GAPDH | GTCTCCTCTGACTTCAACAGCG | ACCACCCTGTTGCTGTAGCCAA | | |
| Actin | CACCATTGGCAATGAGCGGTTC | AGGTCTTTGCGGATGTCCACGT | | |

**Table S1**. List of Antibodies and primers used.
